## Supplemental figures for "Shortcutting from self-motion signals: quantifying trajectories and active sensing in an open maze"

### Supplementary Figures

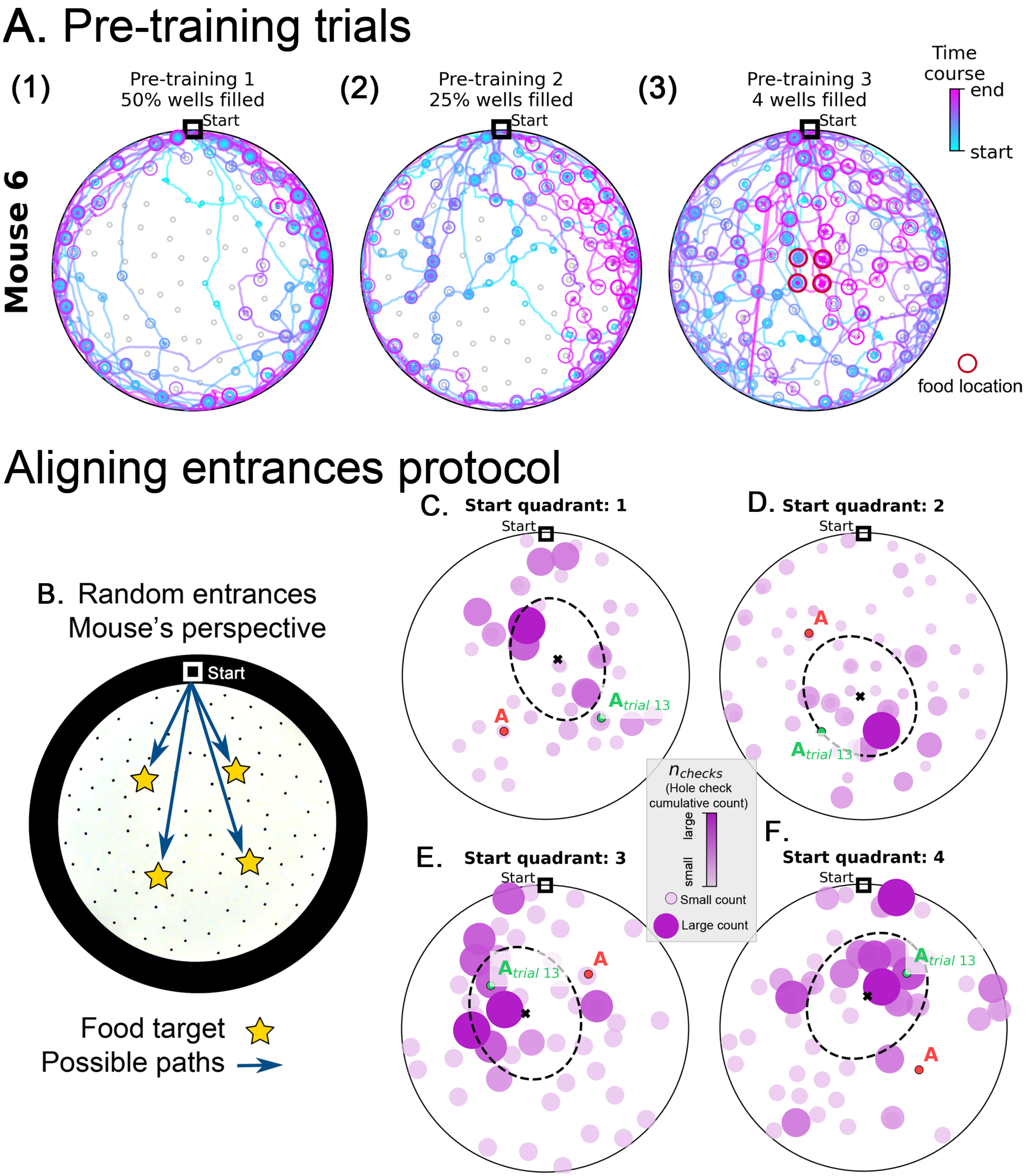


**Figure S1. Pre-training trajectories and Active sensing from the mouse perspective in random entrance experiment**. **Panel A.** Example trajectories of a mouse during pretraining (see Methods). The mice are habituated to their home cage (Day 1) and the maze (Day 2, no food). On Day 3, mice start exploring an arena with randomly selected 50% of the holes filled with food (panel A1). On Day 4 25% of the holes are filled with food (panel A2). Most trajectories are confined near the maze walls on Days 3 and 4. During the final pre-training Day 5, only the four central wells (panel A3, red circles) have food and the mice now explored the entire maze. Circles indicate hole checks (with or without food) with color and size indicating the time course of checking – small, blue circles occur early along the trajectory and pink large circles later in the trajectory. Food locations are only illustrated for Day 5 where they are confined to four central holes. The mice can re-enter their home cage at any time during the food search resulting in multiple blue and magenta lines converging at the Start location. **Panels B-F. Active sensing from the mouse perspective in random entrance experiment. B.** The mouse can enter the arena from each of the four quadrants of the circle. Due to the circular symmetry, we can rotate each entrance setup so that it always aligns to the top of the screen, as shown. This is the mouse perspective of the experiment, where now the mouse has to find one of four possible target locations. Panels **C**-**F** show the hole checks of trial 14 of the Random Entrance experiment as an example (target=red "A" label). The previous spots of the target are shown with a green "$A_{trial\ 13}$" label and depend on the specific entrance a particular mouse took in trial 13. Quadrants are defined in main Fig. 1. Starting from Quadrant 1 is linked to the configuration in **panel C**; quadrant 2 in **panel D**; quadrant 3 in **panel E**; and quadrant 4 in **panel F**. Moreover, notice that the configuration of targets for panel D is rotated $90^o$ clockwise relative to panel C; panel E is $180^o$ clockwise relative to C; and F is $270^o$ clockwise relative to panel C. Thus, we can reverse these rotations and align all panels to the configuration in C (see Experiment Alignment in Methods). We do that consistently for every trial, increasing the sample for the active sensing at each trial. The data shown in panels C to F refer to the frequency of hole checks in the arena: number of checks accumulated over mice divided by total number of checks, normalized between zero (smaller circles colored in light pink) and one (larger circles colored in dark pink). Larger balls represent higher normalized check rates.

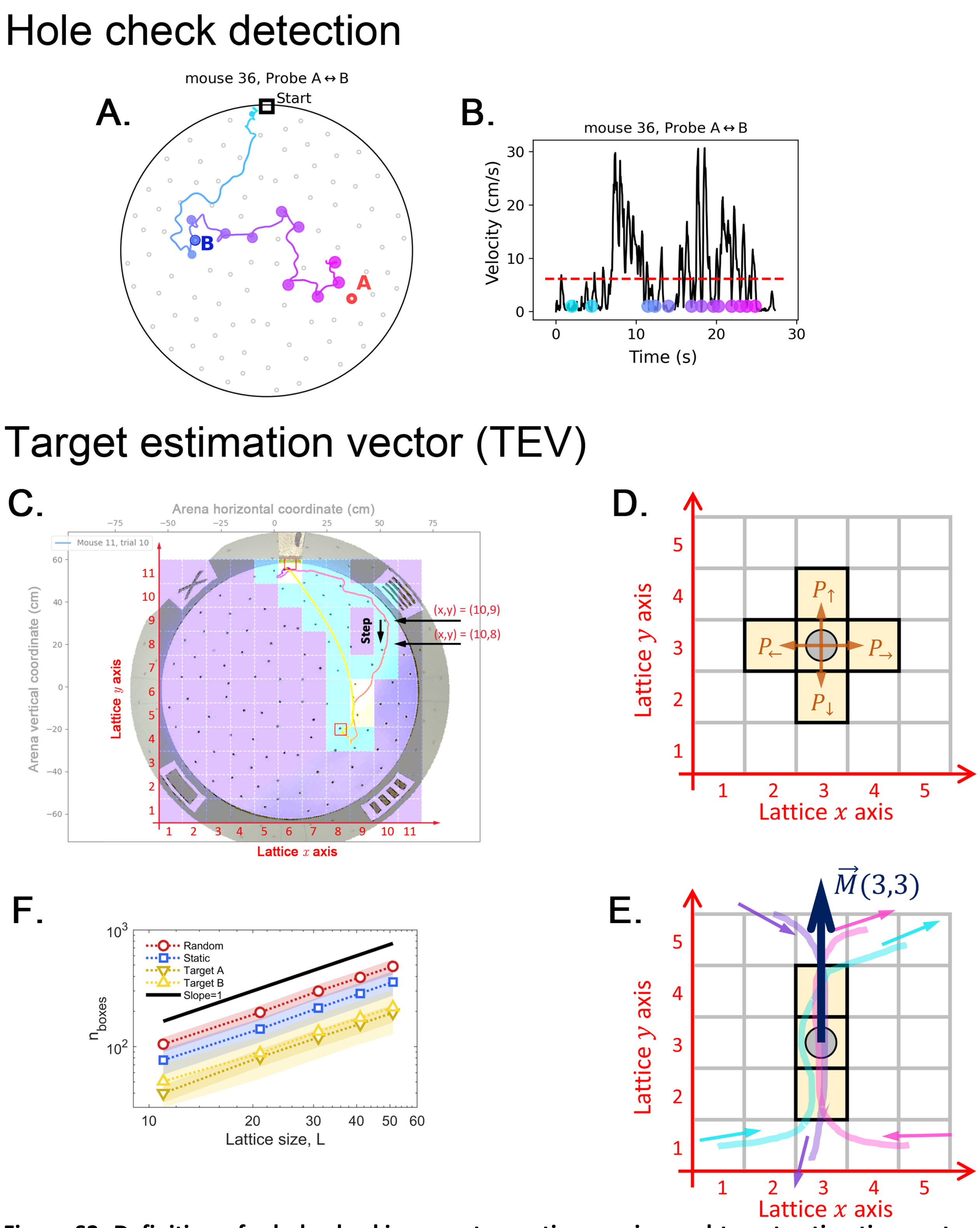


**Figure S2. Definition of a hole-checking event as active sensing and target estimation vector (TEV). A.** Observed trajectory with hole-checking events identified by the two criteria sets of our method, applied subsequently to avoid missed checks. **B.** The velocity profile with dots representing the hole-checking events shown in panel A. The color from cyan to magenta represents the time order of the events, as presented in panel A. See Hole-checking Event Detection in Methods for details of the detection criteria. A video is also available Shortcut video (Video 1) to illustrate the operation of the hole check algorithm and the mouse behavior. **Panels C-E: Arena lattice, box size scaling and step map definition. C.** A square lattice with $L^2$ boxes is overlayed on top of the arena recording to build the step map. There are $L$ boxes along both the $x$ and $y$ axes. $L$ is an odd number, such that the center box is aligned with the center of the arena. For $L = 11$, each lattice site (box) measures approximately 11 cm by 11 cm. This lattice is used to calculate the step maps (see Step Map Calculation in Methods). **D.** To generate the step maps, we start by counting the number of steps a mouse takes between any two adjacent sites in the lattice; for example, when the mouse lies in the box marked by the gray circle, it can choose to take a step up, down, left or right. If no trajectories pass over that site towards a given direction, then that particular direction is assumed to have null probability $P_0 = 1/4$. **E.** An example showing three trajectories that eventually pass by the marked spot, two times going up (cyan and magenta) and one time going down. The step map vector $\vec{M}(3,3)$ is then pointing up. See Methods for the detailed calculation of this vector. **F.** We count the number of boxes $n_{boxes}$ for each trajectory (exemplified in panel a) and plot it versus the lattice lateral size $L$, yielding the box dimension[1] equal 1 for all trajectories in every trial and experimental setup. The observed robust scaling allows us to use moderately small $L = 11$ for our experimental analysis; thus, larger $L$ yields qualitatively the same results.

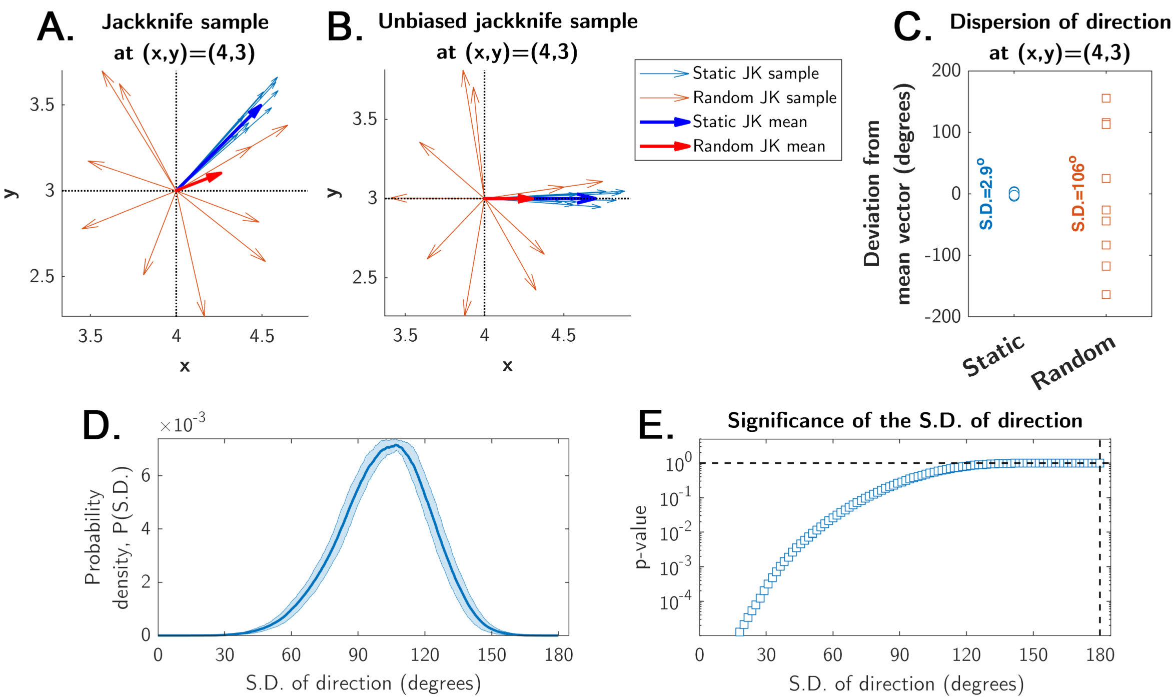


**Figure S3. Estimation of significance for the mean displacement direction calculation.** The jackknife sampling procedure is obtained by the "leave-one-out" rule, yielding N unique jackknife samples of N-1 points from an original sample of N points (see Methods). **A.** Illustration of a jackknife sample with N-1=10 vectors for a single site in the arena for both static (blue, strong directionality) and random (red, weak directionality due to a more uniform direction distribution) entrance experiments (pale-colored arrows=sample; bold-colored arrows=mean; the means make up the displacement maps shown in main Figs. 4,5,7,8, and Fig. S7). **B.** The same sample from panel a, but all vectors' angles (i.e., directions) were shifted such that the mean angle of each condition (static and random entrances) is zero degrees. This procedure does not change the distribution of angles, but makes all angles distributed with zero mean between -180° and 180°. **C.** The angles of the mean displacement vectors of a typical site in the static experiment are narrowly distributed around 0° (S.D.=2.9° for this particular site, after the unbiasing in panel b); the angles of the random entrance experiments are widely distributed (S.D.=106° for this particular site), reflecting weak directionality. **D.** The probability density of observing some particular value for the S.D. of a collection of N=8 uniformly distributed angles between -180° and 180°. It was calculated numerically (shaded area is the error of the estimated probability density function from 10,000 independent realizations). **E.** The p-value represents the significance of a given S.D. shown in panel d (i.e., probability of observing the S.D.); it is the integral of the probability density in panel d up to S.D. (see Methods). This p-value shows that the probability of observing a S.D. < 15° is negligibly small ($p<0.0001$). This means that observing such an S.D. in the static entrance experiments cannot be due to chance. These p-values are the ones shown for the displacement maps in main Figs. 4,5,7,8.

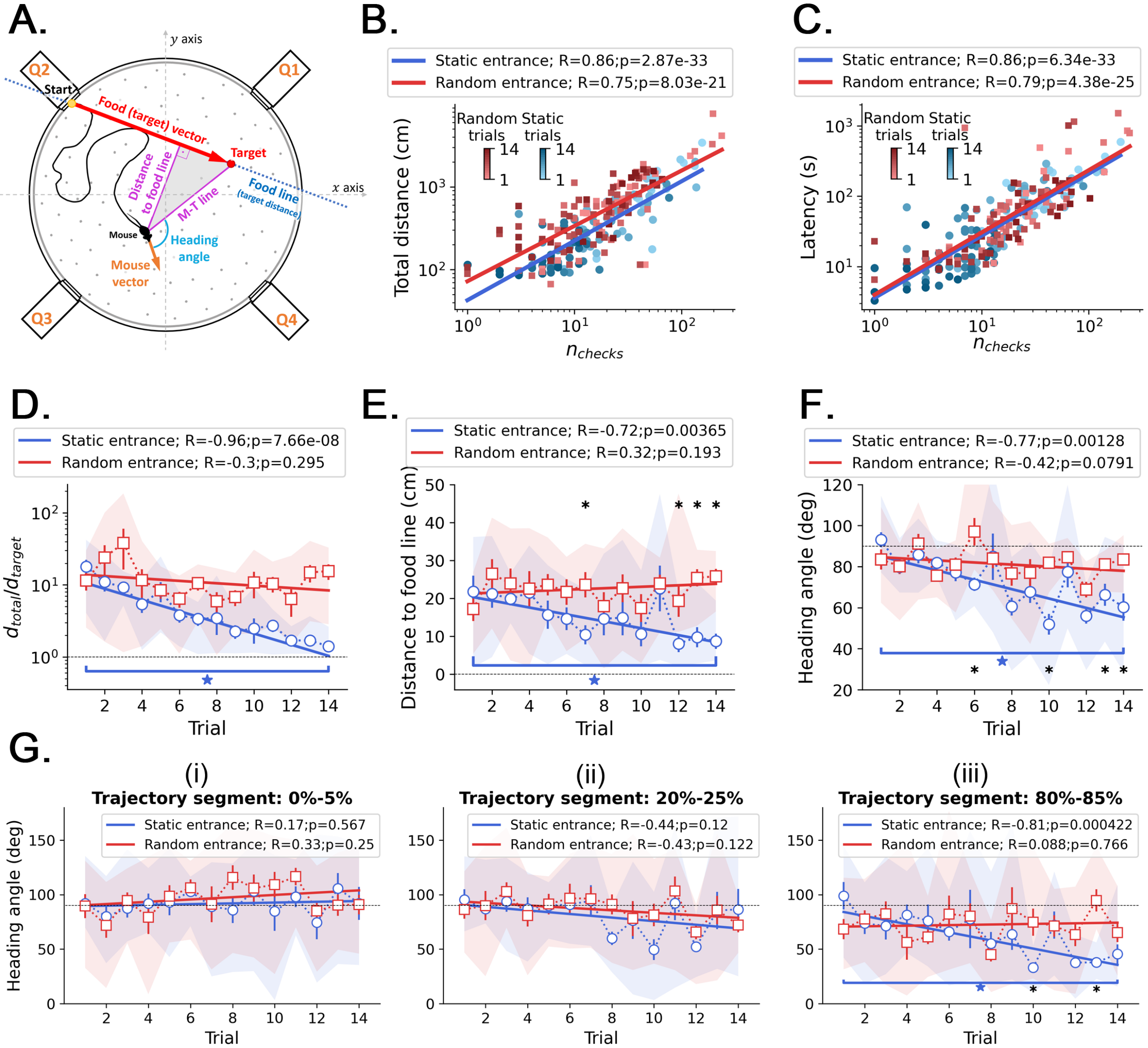


**Figure S4. Kinetic and geometric features and correlation with active sensing. A.** Definition of the geometric features that measure performance across trials. *Food line:* the straight line that connects the food hole (target) to the entrance; this line defines the optimal trajectory and minimum distance from start to target, $d_{target}$. *Distance to food line*: minimum (perpendicular) distance from the mouse center to the food line (it can be regarded as the trajectory "error", or deviation, from the optimal trajectory). *M-T line*: a line that connects the mouse center to the target. *Mouse vector*: a vector that points from the mouse center to its nose. *Heading angle*: angle between the mouse vector and the M-T line (heading angle=0 means going straight to the target). **B.** The total traveled distance $d_{total}$ correlates with the number of checked holes $n_{checks}$ for every trial, sampling all mice together in both random and static entrance experiments. **C.** The total trial time spent before reaching the food almost linearly correlates with the number of checked holes $n_{checks}$ for every trial: regardless of how long the mouse runs, they keep checking the arena for food. This is evidence of path integration because even after learning, the mice are still uncertain, so they keep checking. In the main text and in Fig. S9, we showed that these checks in later trials accumulate near (<20cm) the target. **D,E,F.** Comparison of the evolution of kinetic and geometric parameters across trials, averaging each performance feature over mice (N=8; symbols=mean; shaded region=full extent of the sample; error bars=standard deviation). **D.** The traveled distance $d_{total}$ (**panel D**) becomes quasi-optimal in the static entrance case, approaching $d_{target}$. **E.** The heading angle to the food line decreases significantly only in the static case. **F.** The heading angle decreases significantly only in the static case. Asterisks mark significant differences between random and static cases (q<0.05, FDR-corrected); stars mark significant differences between first and last trials of the same case (p<0.05). **G.** Heading angle averaged over mice displayed as a function of trials for different segments of the trajectory: (i) 0%-5%; (ii) 20%-25%; and (iii) 80%-85%. A segment is defined as the percentage of the latency (i.e., total trial time). For example, the 0%-5% comprises only the part of the trajectory between the start and 5% of the total trial time shown in **panel C**. Only in the last parts of the trajectories the mice turn to the food (evidenced by a significant decrease of the heading angle vs. trials in the 80%-85% segment), suggesting that the mice tend to keep their trajectories variable.

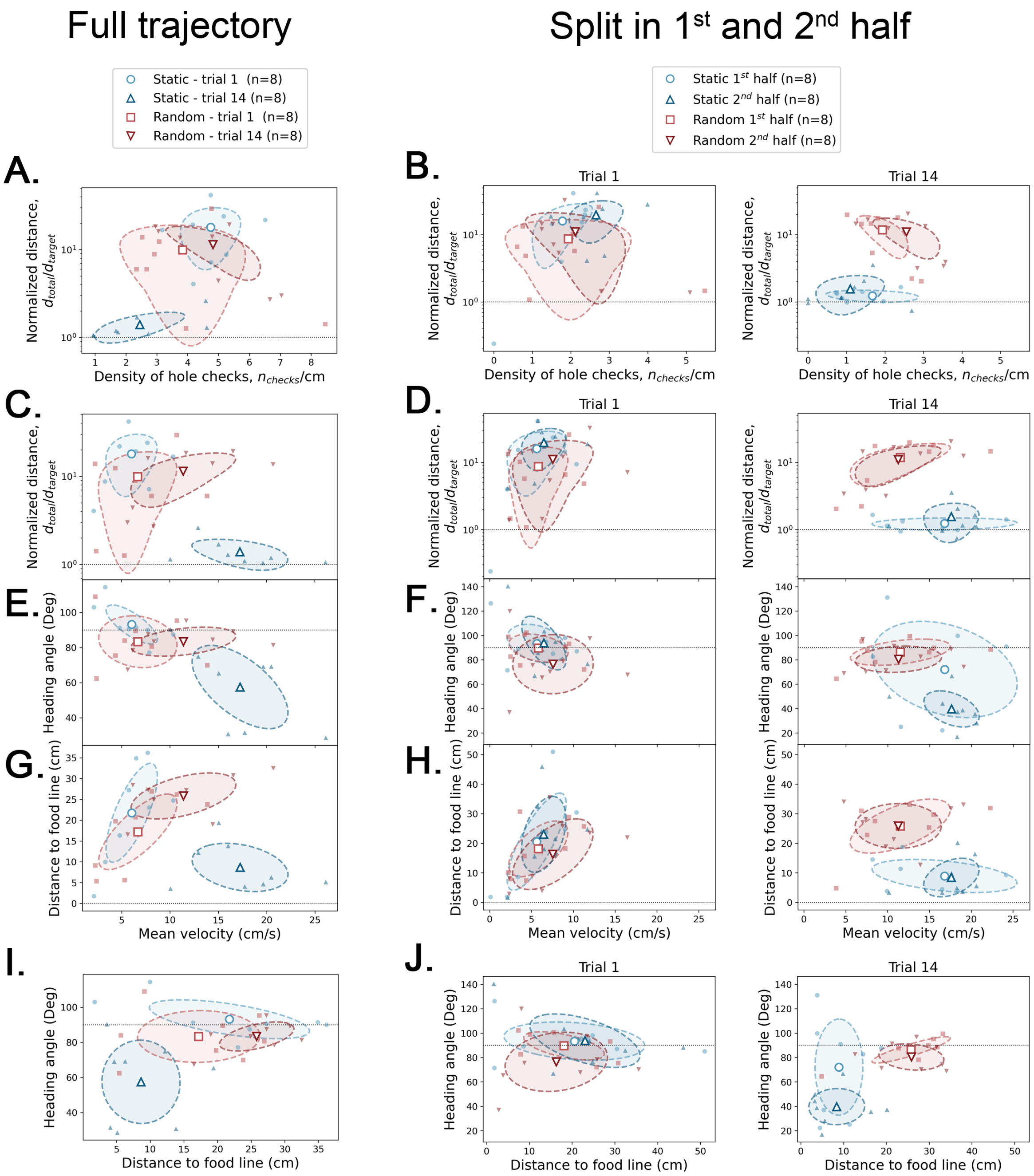


**Figure S5. Covariance between geometric and kinetic features, and active sensing.** The definition of each of these parameters is given in Fig. S4. Small filled symbols=average over trajectory for each mouse; large empty symbols=average over mice (N=8) of small symbols. Dashed ellipsis=covariance across mice. **Left column**: full trajectory analysis – each feature is averaged over the whole trajectory of each mouse. **Center and right columns**: half trajectory analysis – each feature is averaged over the first and second half of the trajectory for trial 1 (center) and trial 14 (right). The four large empty symbols give the average over the eight mice for each case. Horizontal dotted lines are there for reference values.

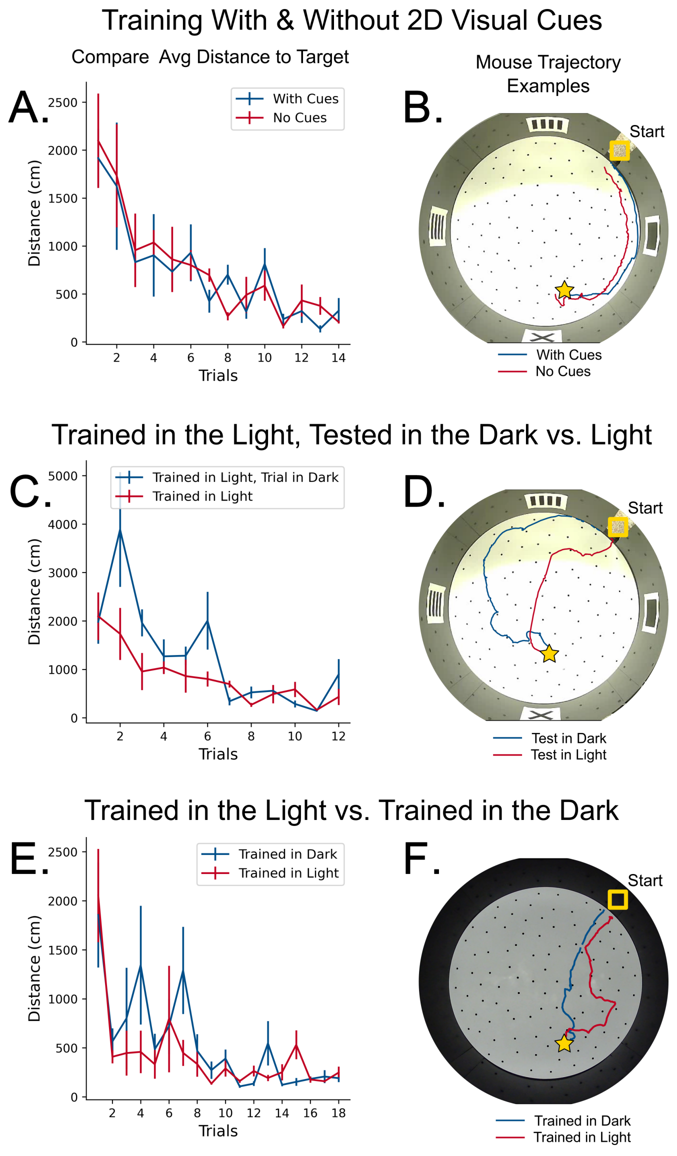


**Figure S6. Control experiments for path-integration. A.** Comparison in learning rate between mice with 2D cues on the wall (blue) and mice with no cues (red). N = 4 mice per group, Error = SE. No significant difference in performance. **B.** Example trajectories of mice trained with and without 2D visual cues. **C.** Comparison in learning rate between two groups of mice trained in the light. N = 4 mice per group, Error = SE. No significant difference in performance. **D.** An example trajectory of a mouse trained in the light successfully navigating towards the target in darkness (blue line). A control trajectory of a mouse navigating in the light is shown (red line). **E.** Comparison in learning rate between mice trained and tested in complete darkness versus mice trained in light and tested in light. The trained and tested in darkness mice had additional controls for potential odor cues (see Methods). N = 4 mice per group, Error = SE. No significant difference in performance. **F.** Example trajectories of mice trained and tested in darkness versus trained and tested in light successfully traveling towards the target in both darkness (blue line) and light (red line) respectively.

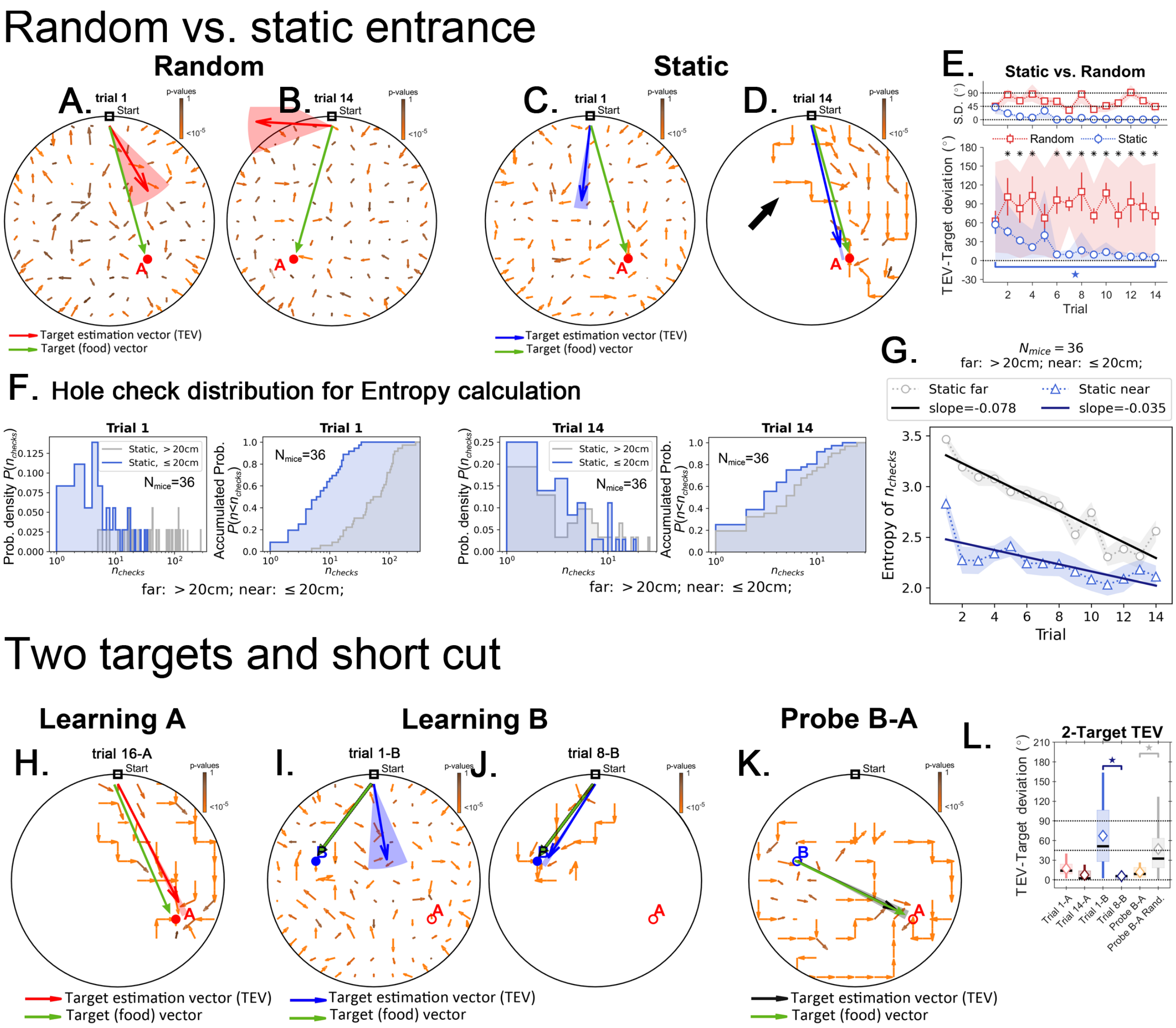


**Figure S7. Target estimation vector (TEV) in detail for the random, static and two-target experiments.** Black arrows point to the mean displacement direction starting from any given site in the arena. **Panels A-D and H-K** green arrows are the target (food) vectors (point from start to target; or from B->A in the Probe B-A trial) and are to be compared with the TEVs. These panels have the same data as in Figures **4 and 7**, respectively, although here all the mean displacement vectors are displayed without censoring large p values. Here, the color of the arrows corresponds to p-values (brown=1; orange=0). In the main text, only displacements with very small p-values are shown (see main text and Methods). **Panels A and B show the random entrance condition.** The TEV (red) reflects the learned target position, and does not correspond with the food vector, highlighting a random search pattern. **Panels C and D show the static entrance condition.** The TEV (blue) becomes closely aligned with the food vector during late training [**panel D**]. The big black arrow highlights how trajectories are still variable even after learning the TEV. The variation observed in trajectories result in a TEV pointing almost directly to the food; error is 11 cm – the size of the lattice site, corresponding roughly to the average distance between nearest holes. **Panel E:** the angle between food vector and TEV in both random and static conditions as training progresses. Angle differences decrease in the static case but not in the random case. N = 8 per group, significant differences are marked with an asterisk (FDR corrected, $q<0.01$). The star marks significant difference between first and last trials only for the static entrance case. **Panel F:** distribution of the number of hole checks, $P(n_{checks})$ near the target (≤20cm) vs. far for all experiments in static entrance experiment (N=36 mice). **Panel G:** entropy related to the distributions in panel F, $H = -\Sigma P(n_{checks})\log[P(n_{checks})]$, where the sum runs over all values of $n_{checks}$. The entropy of the number of hole checks is always lower for the near compared to far hole checks. In other words, the mice are more consistent in choosing the numbers of holes to check when near the food site (<20 cm) versus further from the food (>20 cm). The entropy decays for both far and near the target conditions consistent with learning food location over trials, but the decay is twice as steep for the far condition. The cumulative densities are shown next to the probability densities.

**Panel H.** In the static condition the TEV (red) points to the learned location of the target A (analogous to Static target in **panel D**). **Panels I and J.** First and last training trials of target B (after completing training of target A)**.** The TEV (in blue) starts pointing to A (trial 1B), but ends up pointing to B (trial 8B). **Panel K. No food probe trial.** The mean displacement generates a TEV (orange vector) that points from site B to A, highlighting the shortcut route. **Panel L. Learning vs. shortcut.** The difference in angle between the food vector and the TEV. The difference becomes significantly lower as training progresses, going back up as the target is switched from A to B, and then going down again as the mice learn the new target. During the probe trial, the TEV-target angle is as small as the late learning trials of either target B or target A (i.e., when the mice know their route). The TEV for the probe trial is significantly smaller than a TEV generated using a random step map (gray box, see Methods for how a random step map was generated).

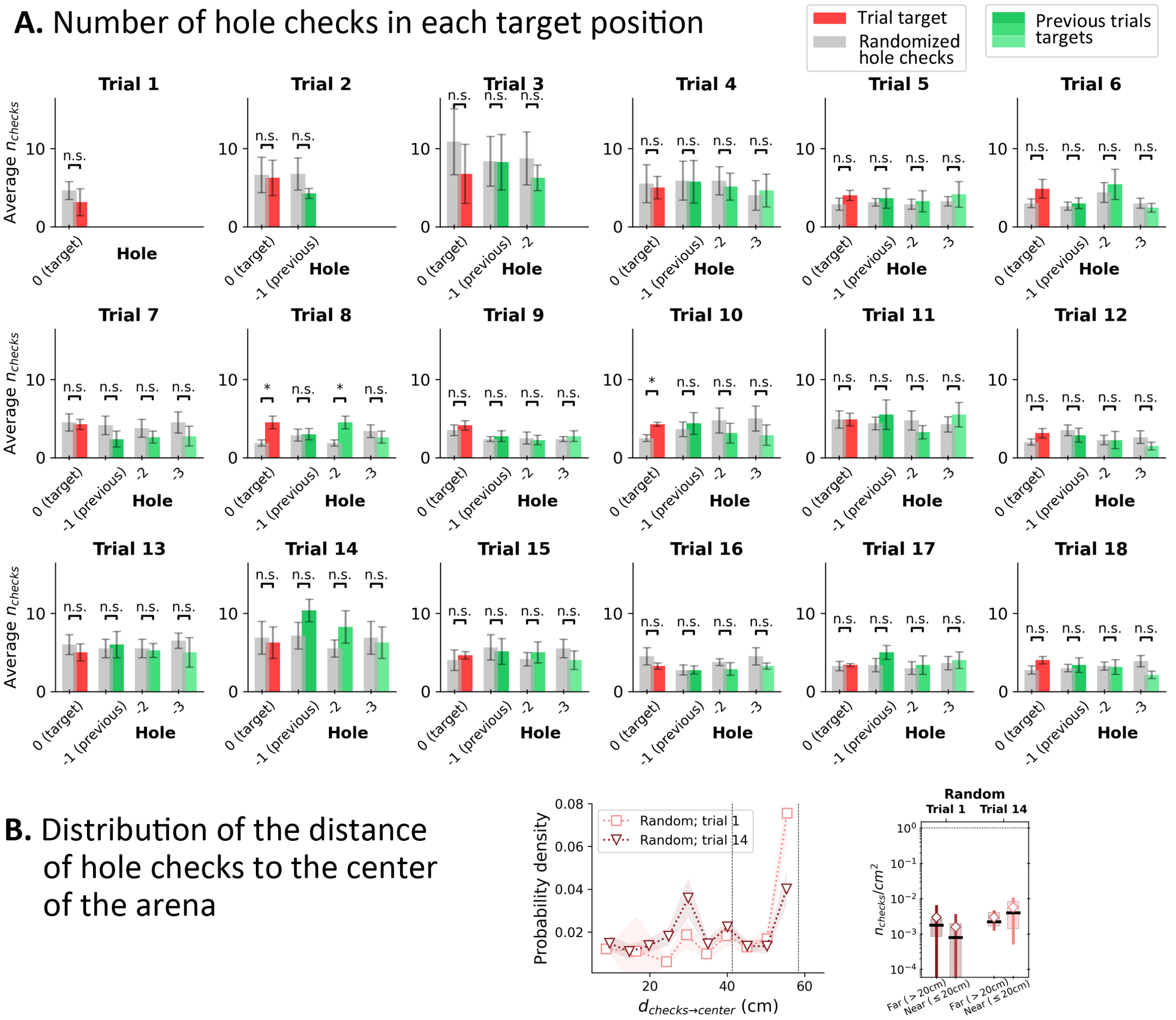


**Figure S8. Hole check distribution in Random Entrance protocol for the four target holes from the mouse's perspective. A.** Red bar: hole checks in the target hole (labeled "0"). Green shaded bars: hole checks in the previous positions of the target in the mouse's perspective (see Fig. S1b; "-1" for the position of the target in the previous trial; "-2" for the trial before previous; and "-3" for the trial before "-2"). Since, there are training trials, each of the target holes (0, -1, -2 and -3) had food in their respective trial (i.e., "0" has food in the trial; -1 had food in the previous trial; -2 had food in the trial before previous; and -3 had food in the trial before -2). Gray bars: randomized hole checks for each corresponding position: all detected hole checks in a given trial were uniformly distributed across the 100 holes of the arena, and then we counted the number of checks in each of the four targets' positions. Whiskers: standard deviation (n=8). Gray bars were compared to the corresponding colored bars via a paired t-test. Significant differences were rarely found (Trial 8 only) between randomized data and actual hole checks, suggesting that the mice are performing randomly. n.s. = not significant. **B Left:** Distribution of the distance of the hole checks to the center of the arena (to be compared with Fig. 2I and Fig. 3I in the main text). The distance of the active sensing to the center of the arena is almost uniform, indicating that the mice have no preferential distance to check for food. **B Right.** The density of checks near the center (<20cm away) is the same as farther away. This suggests that there is no obvious spatial learning feature relating the target with the central spot of the maze.

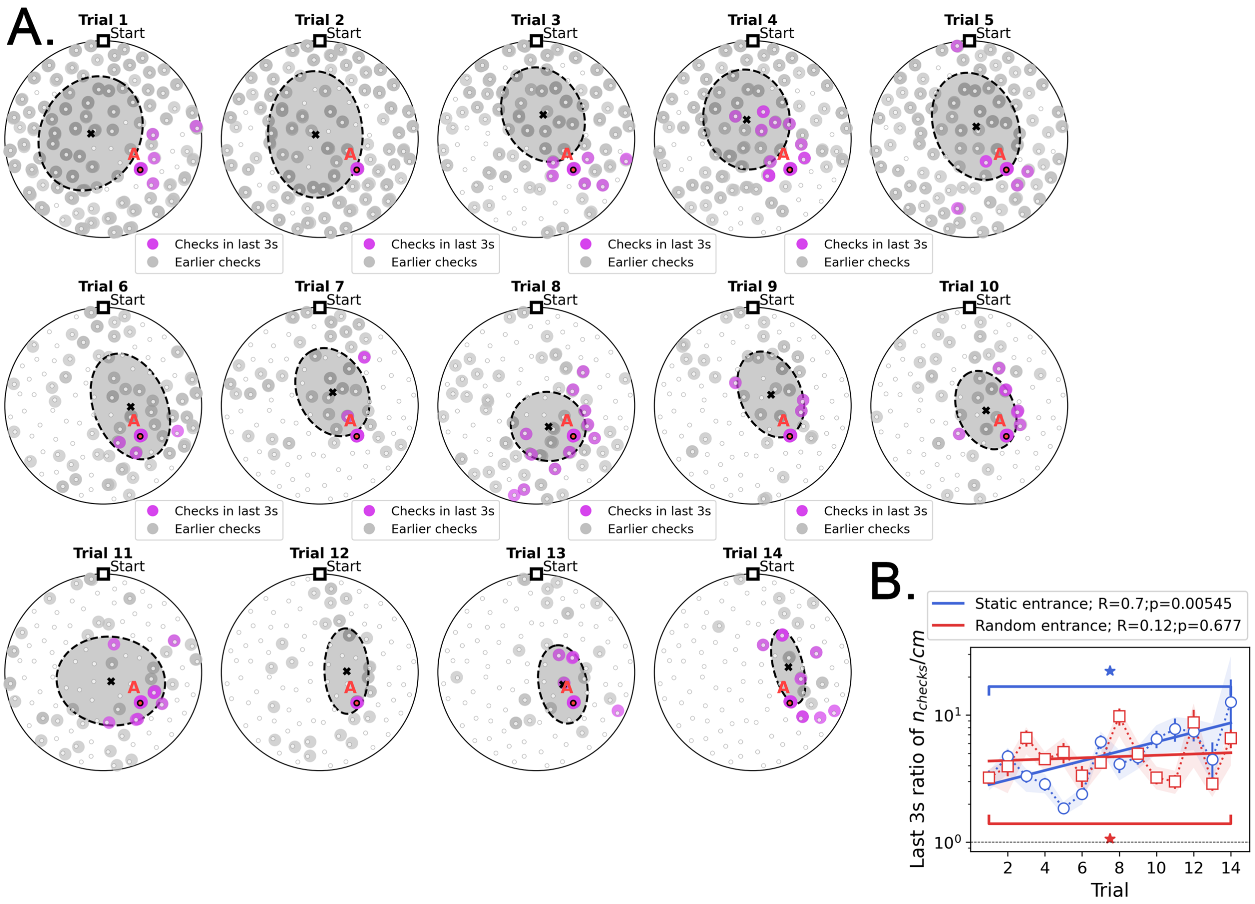


**Figure S9. Location of hole checks in the last 3 seconds before finding the target in Static Entrance protocol. A.** Pink spots: hole checks in the last 3s before finding the target; gray spots: earlier hole checks. Dashed ellipsis (x=mean): dispersion (covariance) of the spatial distribution of hole checks. The distribution of hole checks migrates towards the target (labeled A) as the mice learn. All mice (n=8) pooled together. **B.** Density of hole checks ($n_{checks}$ per traveled distance) in the last 3s (pink spots), compared with that density for the earlier path (gray spots), as a function of trial. There is significant increase over trials (R=0.7, p=0.00545) for the static entrance case, implying that the hole checks are converging to be temporally "close" to the food as the mice learn – in the first few trials, holes are checked throughout the entire duration of the run. We hypothesize that the pink spots might act as anchors for place fields. Shade behind the curve marks the full extent of the sample (n=8 mice). Stars: significant difference between first and last trial (paired t-test, p<0.05).

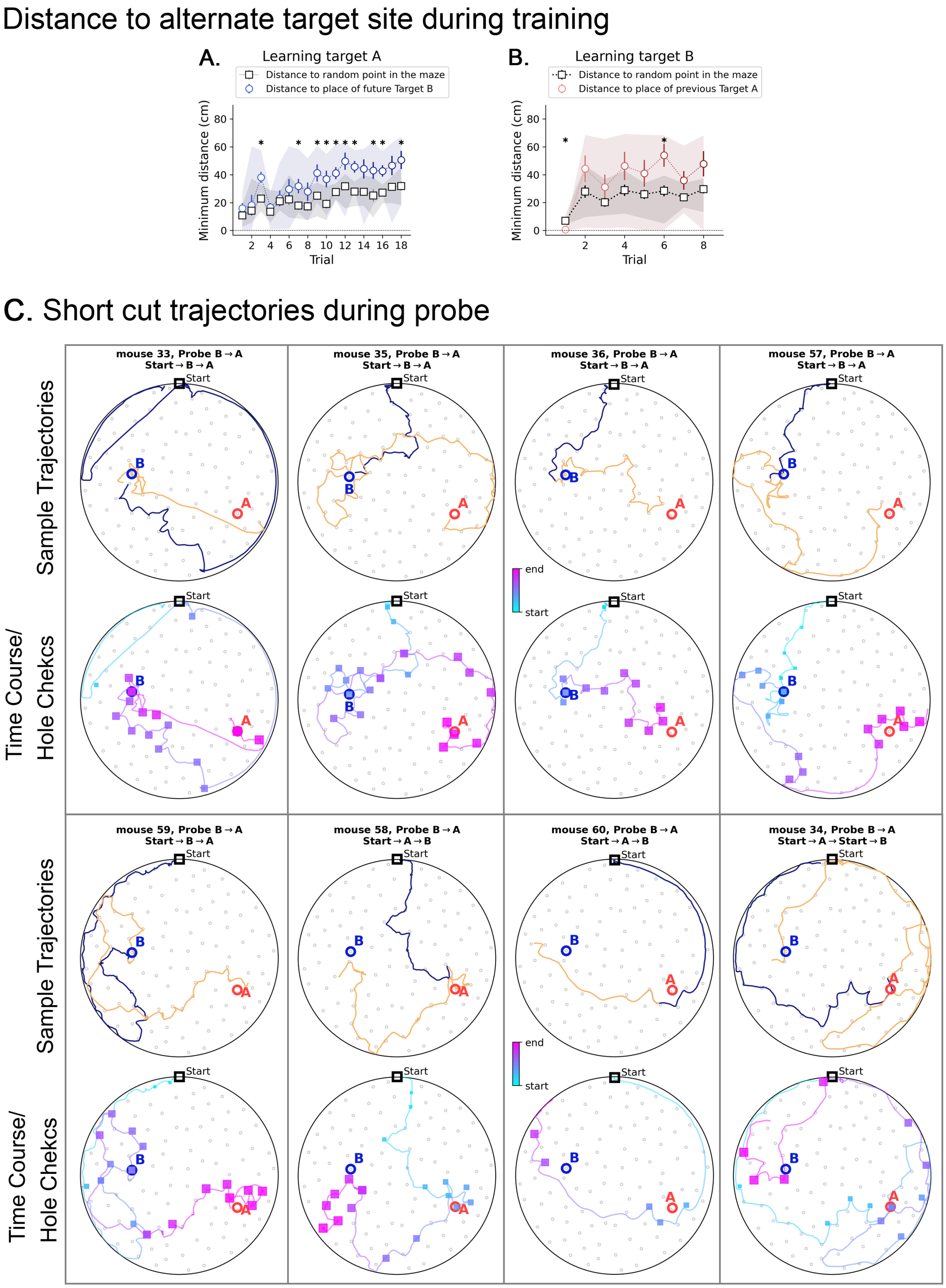


**Figure S10. Minimum distance to alternative target in 2-target condition, and short cut trajectories. A and B.** Black squares are the minimum distances expected by chance for each trial; they are calculated with respect to a random point, and then averaged over ten such random points. Shaded area represents the range of values, and bars represent the standard deviation.

**A.** During the learning of target A (no previous target has been presented), the mice keep consistently farther than expected by chance from the location where Target B will be placed (blue circles; asterisks show significant differences to chance). **B.** The same pattern repeats when learning target B (target A has been already learned and is not present anymore): the mice keep consistently farther than expected by chance from the location where the previous Target A was found (red circles; asterisks show significant differences to chance). During the learning of B, some mice did visit the A location sporadically, making the shaded area larger in panel **B**. However, they did not rely on target A to find B.

**C: Short cuts and active sensing for the Probe** B→A **trial.** For each mouse, the plots in the top arenas contain trajectories split into two parts: before (blue) and after (orange) the visit to the first target (either A or B, whichever comes first, colored in blue). The bottom arenas of each mouse contain the time course (blue is early and pink is late) with hole checks in squares (blue is early and pink is late) to give a sense of when each event happened. We cut off the trajectories after visit to the last target (either A or B, whichever comes later, to ensure we observe its visit to both A and B locations). The orange trajectories are the ones that the mice took after realizing there is no food in either A or B (whichever comes earlier). We expect that mice perform the sequence Start→B→A, since B was trained last. Notice in the time course plots that mice 33, 35, 36, 57 and 59 do exactly this without ever going back to Start. Mice 34, 58 and 60 perform Start→A→B, but only mouse 34 visits the Start before heading from A to B.
